# A breeder in the sky: scoring flowering with fewer flights

**DOI:** 10.64898/2026.09.19.752900

**Authors:** Zhongjie Ji, Aaron J. DeSalvio, Alper Adak, Mustafa A. Arik, David Ertl, Shawn M. Kaeppler, Jode Edwards, Sherry Flint-Garcia, Candice N. Hirsch, Addie M. Thompson, Jacob D. Washburn, Natalia de Leon, Bijesh Maharjan, Dipak Santra, Pascal Izere, Biquan Zhao, Yeyin Shi, Seth C. Murray, Yufeng Ge, James C. Schnable

## Abstract

Flowering time is a trait of broad interest and importance to both plant breeders and plant biologists. Unlike many other traits, flowering time cannot be scored accurately from a single observation; its measurement requires repeated observations of the same experiments over time. Both classification- and object-detection-based approaches have demonstrated the potential to estimate flowering time from Unmanned aerial vehicle (UAV) imagery rather than by manual observation, but they still require data collection at many time points. Here, we train and deploy a regression-based framework for scoring flowering time. Using a dataset of more than 200,000 UAV images and associated flowering-time records collected across 27 environments, we demonstrate that this approach enables the prediction of flowering time from as few as one observation per field experiment and successfully generalizes to field experiments in environments not represented in the training dataset. These results indicate that regression-based approaches for predicting flowering time from sparse UAV observations have the potential to substantially reduce the data-collection burden in field experiments.

## INTRODUCTION

Flowering time plays a critical role in the suitability of crop cultivars for specific regions and in the adaptation of wild plant accessions to ecological niches. Because a large proportion of variation in this trait is attributable to genetics in many species, and because the trait affects both crop productivity and plant fitness (Tollenaar et al., 1979; Bonhomme et al., 1994; Žalud et al., 2017), flowering time is often used as a model trait in quantitative- and population-genetics research (Buckler et al., 2009; Weis et al., 2014). Changes in flowering time can produce substantial differences in many other plant phenotypes (Cross and Zuber, 1973; Cui et al., 2017; Li et al., 2016; Borrás et al., 2009; Ji et al., 2025); therefore, researchers who are not studying flowering time directly must often score this trait to control for its confounding effects. Finally, plant breeders record flowering time early in the breeding process as a necessary step in identifying varieties that may be suitable for different maturity zones.

However, unlike many other plant traits (e.g., height, grain yield, leaf number, and root architecture), variation in flowering time within a field experiment cannot be measured on a single date. Instead, repeated visits are necessary from the time the earliest individual flowers until the latest individual flowers to determine when each plot or accession reaches a defined flowering threshold. The need to observe each plot every 1–3 days during the flowering season to obtain accurate data (Pauli et al., 2016) has historically imposed a major labor burden on plant breeders and plant geneticists.

In the past decade, unmanned aerial vehicles (UAVs) have transformed field phenotyping by offering fast, repeatable, and high-resolution monitoring at scale. Equipped with RGB cameras and spectral sensors, UAVs can capture fine-grained information on crop growth, health, and architecture (Jin and Eklundh, 2014; Vidican et al., 2023; Chang et al., 2020). Several approaches have been used to estimate flowering phenology from image and environmental data. Models trained to distinguish plots that have or have not flowered have been applied to UAV-collected RGB images (Shepard et al., 2025), while related image-analysis approaches have quantified visible floral structures using pixel-based classification, clustering, and density-based counting methods (Vanbrabant et al., 2020; Li et al., 2023; Zhao et al., 2022). Classification-based frameworks have also been used to predict flowering time from genotype and environmental time-series data (Deva et al., 2024). However, although these approaches reduce the need for repeated manual field scoring, they still require frequent UAV data collection and image processing throughout the flowering period. Alternative approaches using multispectral imagery have demonstrated the potential to predict flowering time from a small number of flights across the window of flowering in a field experiment, but currently exhibit poor generalization to new locations or years (Camenzind and Yu, 2024; Fan et al., 2022).

Here, we develop and test an approach for measuring flowering time that redefines the task from a classification or object-detection problem to a continuous regression problem. Instead of classifying images as flowering or non-flowering, or detecting specific visible reproductive structures, we assign each UAV image of a plot a quantitative label representing the temporal distance between image collection and flowering for that plot. This approach allows the model to identify and use information about phenological transitions both before and after flowering rather than relying solely on the flowering event itself. We demonstrate that this approach can score flowering time without requiring many flights throughout the flowering window, that the regression framework can integrate multilocation, multiseason UAV datasets in which individual flights occurred on different dates relative to peak flowering, and that models trained and evaluated on more than 200,000 plot-level images of maize plants with associated flowering-time records can predict flowering time in new experiments and environments.

## MATERIALS AND METHODS

### Data sources

The flowering-time dataset used in this project consists of data from 27 unique environments in which hybrid field trials were conducted, UAV images were collected, and flowering time was scored as part of either the Genomes to Fields (G2F) project or the USDA/NIFA High-Intensity Phenotyping Sites program (HIPS) (Figure 1 and Table S1). The dataset spans seven major corn-growing states across multiple years, with more than 1,200 hybrid genotypes represented in one or more of the included field trials. Environments were defined as unique combinations of year, location, and management practice (e.g., irrigated or nonirrigated and early or late planting). Field-observed flowering time was recorded on a plot basis using the standardized protocols defined by the Genomes to Fields consortium (Pauli et al., 2016; AlKhalifah et al., 2018; Lima et al., 2023). Anthesis date was defined as the first day on which half of the plants in a plot had begun shedding pollen from more than half of the main tassel spike. Silking date was defined as the first day on which half of the plants in a plot exhibited visible silks. Field personnel collected these data by walking through the fields every second or third day at each experimental location. UAV imagery was collected by different research groups following different research protocols and using a range of UAV platforms, cameras, and flight heights. These heterogeneous acquisition conditions reflect the multi-institutional nature of the dataset and allowed us to evaluate model performance across varying imaging configurations. Individual environments included between 3 and 25 UAV flights within an analysis window spanning from one month before to one month after flowering. Orthomosaic images were generated from individual UAV-acquired aerial images using either Pix4Dmapper (Pix4D SA, nd) or Agisoft Metashape (Agisoft LLC, nd) as described by DeSalvio et al. (2026), depending on the dataset and data provider. Images representing individual plots on individual dates were extracted from the orthomosaics using polygon shapefiles and a custom Python workflow. The raw resolution of each plot image varied substantially, ranging from 320 *×* 90 to 1550 *×* 390 pixels, but all images were resized to 224 *×* 224 pixels before the analyses described below.

**Figure 1.**
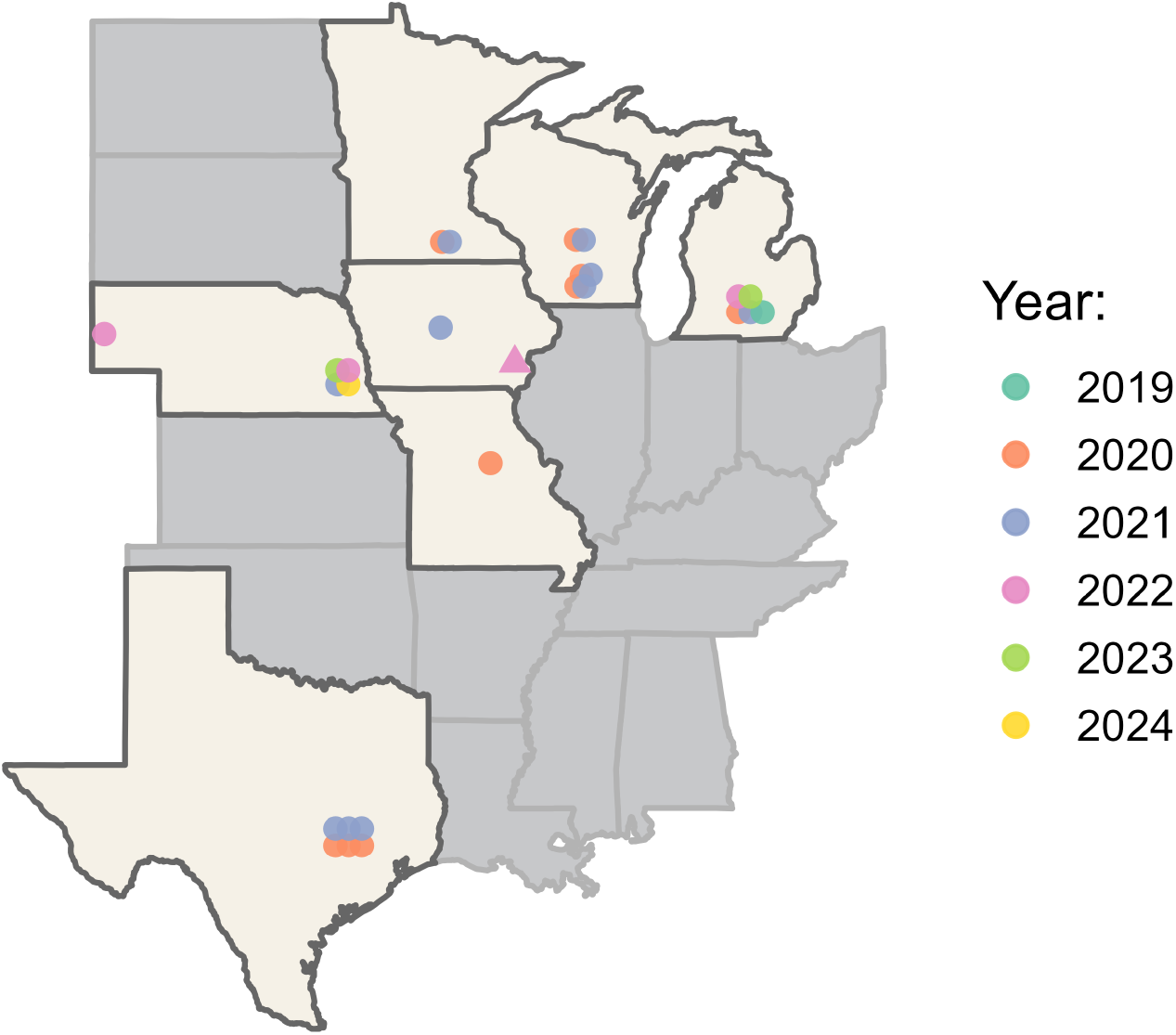
Geographic distribution of UAV data-collection sites used in this study. Field experiments were conducted across seven U.S. states between 2019 and 2024. Circles indicate environments used for model development and evaluation, with colors denoting the year of data acquisition. The triangle indicates the independent 2022 Crawfordsville HIPS site, for which no field-recorded flowering-time data were available and which was used exclusively for external validation.

### Calculation of growing-degree-day flowering times

Plant growth and development rates respond to changes in temperature. In maize, one of the most widely used approaches for accounting for the effect of temperature on developmental rate is to express time in growing degree days (GDD) (Gilmore Jr and Rogers, 1958; Cross and Zuber, 1972). Weather data were retrieved from the Daymet database (Thornton and Devarakonda, 2024) using planting dates and GPS coordinates for all unique environments except Lincoln in 2024, for which Daymet data were unavailable. For that environment, weather records from LNK, the airport nearest the field, were used instead. For the purposes of our study, daily GDD was calculated using the following formula:

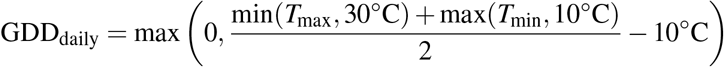

where *T*_max_ and *T*_min_ represent the daily maximum and minimum temperatures (°C), respectively. The base temperature for maize growth was set to 10°C, with temperatures capped at 30°C (Gilmore Jr and Rogers, 1958). The total GDD accumulated over a period of *n* days is given by

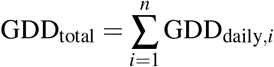

### Formulation of flowering time scoring as a regression problem

To implement this labeling strategy, we developed a continuous flowering label that captures the temporal relationship between each UAV flight date and the observed flowering date for each plot. Specifically, the label was defined as the time difference—measured in either calendar days or GDD—between the image-acquisition date and the field-recorded flowering date. Each UAV survey thus contributed a unique training instance per plot, and the total number of training samples increased proportionally with the number of flights (Figure 2). The flight dates were automatically recorded by the UAV control system, ensuring precise temporal alignment across all experiments.

**Figure 2.**
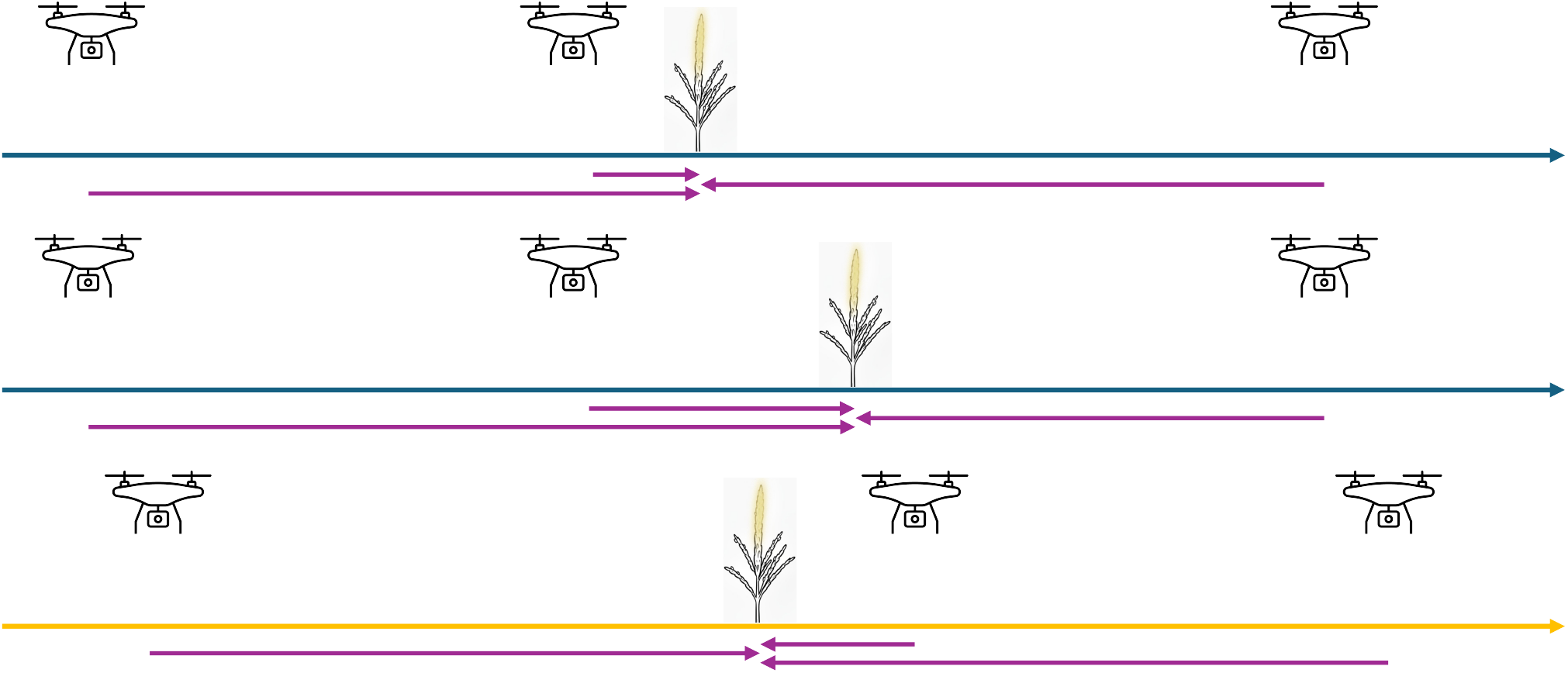
Conceptual labeling framework for using UAV-survey data to predict flowering time. The temporal distance between a UAV survey (UAV icon) and plant anthesis (central plant icon) is used as a predictive label. The upper two panels (dark-blue arrows) show two plots from a single UAV survey with unique anthesis times, whereas the bottom panel (yellow arrow) represents a different experiment. The bidirectional purple arrows indicate the relative time difference, which is normalized and used as a consistent reference label for all data.

By treating each flight date as an independent observation of plot phenology, this formulation allowed the model to learn flowering progression patterns along a continuous time axis, encompassing both pre- and post-flowering stages. We intentionally defined model applicability in terms of temporal distance from flowering rather than discrete developmental stages, avoiding dependence on stage annotations that are not consistently available or directly comparable across environments. Moreover, this approach substantially expanded the training dataset without requiring additional manual labeling, resulting in more than 200,000 plot-by-flight image observations before data augmentation.

To evaluate how the temporal range of image data affects prediction performance, we trained and tested models using two distinct time windows centered around the flowering event. In the first setting, we used all UAV images captured within approximately one month before and after the flowering date. In the second, narrower setting, we restricted both training and testing to images collected within a two-week window on either side of flowering, thereby reducing the dataset size.

Furthermore, to investigate the impact of temporal label units on prediction performance, we trained separate models using flowering labels expressed in calendar days and in GDD, respectively. This comparison enabled us to assess the trade-offs between training volume, temporal proximity to flowering, and the biological relevance of the time metrics in model performance.

### Implementation and training of the prediction model

We employed the well-established Vision Transformer (ViT-Base/16) architecture (Dosovitskiy et al., 2020) as the backbone for flowering-time prediction. ViTs divide images into fixed-size patches and use self-attention to integrate visual information across the image. The model was initialized with ImageNet-pretrained weights to leverage general-purpose visual features learned from large-scale natural image data (Zhuang et al., 2020). To adapt the model for regression tasks, the original classification head was replaced with a custom regression head consisting of a dropout layer followed by a fully connected linear layer that outputs a single continuous value (Figure 3). This modification enabled the model to predict flowering time in either calendar days or growing degree days (GDD). The entire model, including both the ViT backbone and the newly added regression head, was fine-tuned end-to-end on our labeled dataset. This fine-tuning allowed the model to adapt its feature representations to the domain-specific plant imagery, while retaining the benefits of transfer learning. Dropout regularization was applied to mitigate overfitting during training (Srivastava et al., 2014).

**Figure 3.**
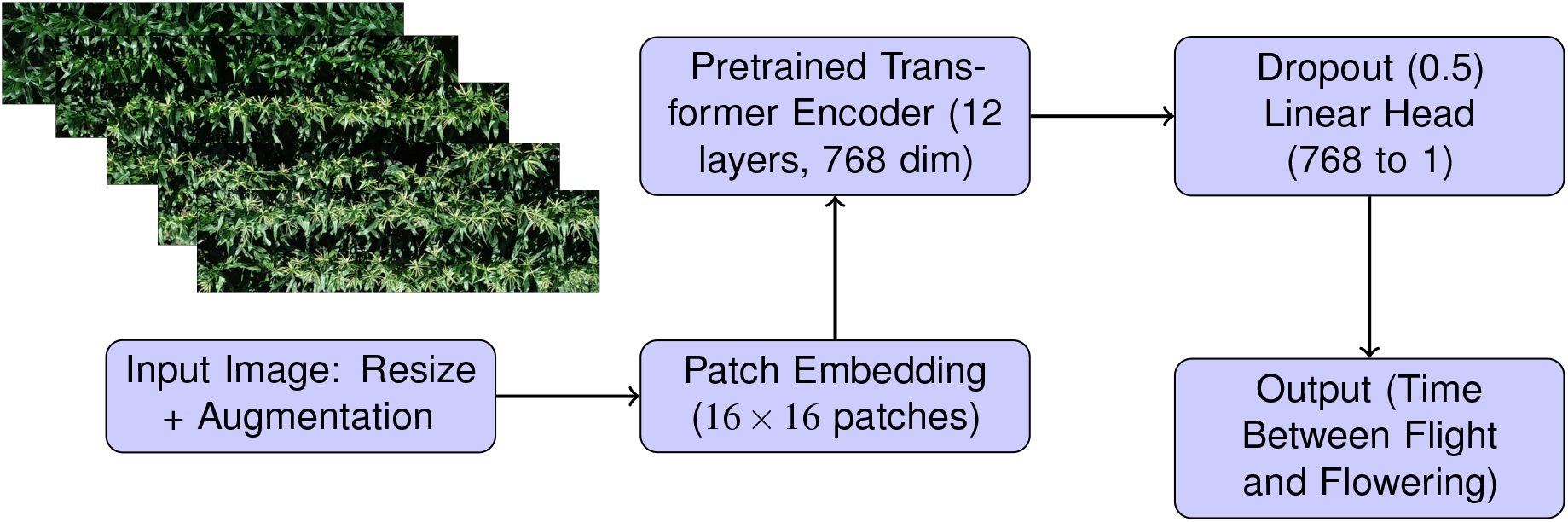
Workflow of the flowering-time regression model. The diagram shows the key processing steps from an input UAV image to the predicted time between the flight and flowering.

We applied a series of data-augmentation techniques before model training. The augmentation pipeline consisted of random horizontal flipping (probability = 0.5), random vertical flipping (probability = 0.2), and random rotations within *±*10°. Additionally, we employed color jittering to increase robustness to variation in illumination and color characteristics across imaging conditions by randomly adjusting brightness, contrast, saturation, and hue. To account for potential imaging noise, we applied a Gaussian blur with a kernel size of 3 and a sigma range of 0.1 to 2.0. All images were converted to tensor format and normalized using the ImageNet mean and standard deviation. These augmentations were implemented using the torchvision.transforms module in PyTorch.

For model training, we used a batch size of 128 images and adopted a mixed-precision strategy to accelerate computation and optimize GPU memory use. The AdamW optimizer was used, with Smooth L1 loss (*β* = 1.0) as the training objective. Smooth L1 loss applies a quadratic penalty to small residuals and a linear penalty to larger residuals, reducing sensitivity to large errors compared with mean squared error.

Model performance was evaluated using mean absolute error (MAE), calculated as follows:

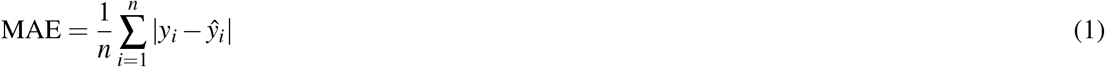

where *y*_*i*_ and *ŷ*_*i*_ denote the observed and predicted values, respectively, for the *i*th sample. Different learning rates were applied for GDD-and calendar-day-based labels: 5 *×* 10^*−*5^ and 1 *×* 10^*−*5^, respectively.

The training schedule included a 10-epoch warm-up phase with gradually increasing learning rates, followed by cosine annealing in subsequent epochs. To mitigate overfitting, an early-stopping strategy with a patience of 10 epochs was implemented, and the model weights corresponding to the lowest validation loss were retained for final prediction.

### Testing and validation of flowering time predictions

Environments were defined as unique combinations of year, location, and management treatment. To prevent data leakage and account for the shared spatial, temporal, and experimental structure of observations within the same environment, we partitioned the dataset at the environment level into training, validation, and test sets using a fixed random seed (42), with approximately 80%, 10%, and 10% of environments assigned to each set, respectively (Roberts et al., 2017). The split was determined before model training and was not based on model performance. This fixed held-out test set was designed to evaluate model performance in previously unseen environments. The test set consisted of 2020WIH2, 2021NEH1, and 2021MIH. For analyses using multiple UAV flights, the model was applied independently to each flight, and the resulting flowering-time estimates for the same plot were combined by taking their arithmetic mean. No multi-temporal model or joint multi-flight input was used.

Repeatability was primarily estimated for genotypes represented in both biological replicates within an environment. Replicate was treated as a fixed effect and genotype as a random effect, with variance components estimated by restricted maximum likelihood. Repeatability was calculated as:

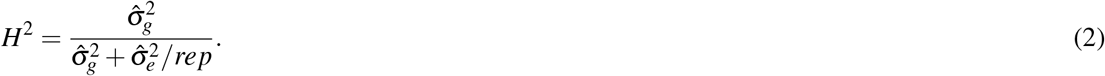

For genotypes represented by multiple plots within the same replicate, plot-level values were first averaged within replicate before estimating repeatability.

Because a substantial number of genotypes in the held-out test environments were represented in only one biological replicate, we additionally calculated Cullis generalized repeatability, which allows information from unbalanced replication to contribute to the estimate:

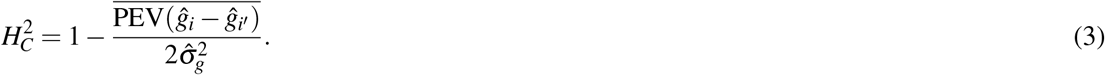

Here, the numerator is the mean prediction error variance of pairwise genotype differences. Cullis repeatability was used as a complementary analysis and is reported in the Supplementary Information. For analyses using multiple UAV flights, repeatability estimates were calculated for each *k*-flight combination and then averaged across combinations. Manual repeatability was calculated from field-recorded flowering phenotypes using a plot set matched to the UAV data. Specifically, we used the subset of physical plots represented in the UAV flight with the greatest plot coverage, with each physical plot included only once.

To evaluate the performance and temporal consistency of our maize flowering-time prediction model without field-recorded anthesis dates, we conducted an independent test using UAV imagery from the 2022 Crawfordsville HIPS site. At this site, 84 maize hybrids were represented across 522 plots, and three UAV surveys were conducted during the growing season (Shrestha et al., 2024). The first two surveys were used for the independent evaluation described here. Because field-recorded anthesis dates were unavailable, each plot in the first orthomosaic was visually annotated as tasseling or non-tasseling based on visible tassels. These annotations served as a surrogate reference. The model predicts the relative time to anthesis for each plot, with negative values indicating pre-anthesis and positive values indicating post-anthesis. For evaluation, predictions were classified as tasseling or non-tasseling based on their sign and compared with the visual annotations. To assess temporal consistency, predictions from the two flights, conducted 14 days apart, were compared for plots that had not flowered by the first flight. The model was trained using imagery collected within two weeks of anthesis, based on preliminary analyses that identified this window as the most informative.

### Visualization of image saliency

To interpret predictions made by the ViT-based regression model, we employed gradient-weighted class activation mapping (Grad-CAM) to visualize salient regions of the input plant images (Selvaraju et al., 2017). Because ViT lacks convolutional layers, we applied Grad-CAM to the final transformer block, specifically targeting the output of the last attention-based feature-projection layer. We used the implementation provided by the pytorch-gradcam library, which supports transformer-based architectures. The resulting activation maps were overlaid on the original images to highlight regions that contributed most strongly to flowering-time predictions. This qualitative visualization helped verify that the model attended to biologically meaningful structures, such as leaf arrangements and canopy patterns, during inference.

## RESULTS

We assembled paired flowering-time and UAV-image datasets collected over a six-year period, from 2019 to 2024, by research groups in seven states (Figure 1). In each environment, anthesis (male flowering) and silking (female flowering) dates were scored and recorded separately for each plot. Individual environments included from 3 to 25 flights throughout the growing season. The resulting dataset included more than 200,000 images of 20,386 unique maize plots. Each image was annotated with the time between image collection and anthesis or silking, expressed in both calendar days and GDD.

Models were trained on data from 21 of the 27 environments. When only images taken within one month of the recorded flowering dates were considered, the models achieved MAEs of 4.92 days (*R*^2^ = 0.85) and 69.17 GDD (*R*^2^ = 0.80) for anthesis, and 4.40 days (*R*^2^ = 0.89) and 58.76 GDD (*R*^2^ = 0.86) for silking, when evaluated on UAV-image data from environments that had been entirely withheld from training (Figure 4). Constraining image collection to within two weeks of flowering improved MAE to 3.20 days (*R*^2^ = 0.79) and 37.17 GDD (*R*^2^ = 0.82) for anthesis, and 3.32 days (*R*^2^ = 0.78) and 40.47 GDD (*R*^2^ = 0.78) for silking (Figure 5). The *R*^2^ values of GDD-based models were consistently higher than those of calendar-day-based models, potentially because GDD more accurately reflects the physiological development observed in UAV images from different sites. MAE was also lower for GDD-based models after converting GDD to calendar days using the typical rate of GDD accumulation at flowering in these test locations (approximately 16 GDD per day).

**Figure 4.**
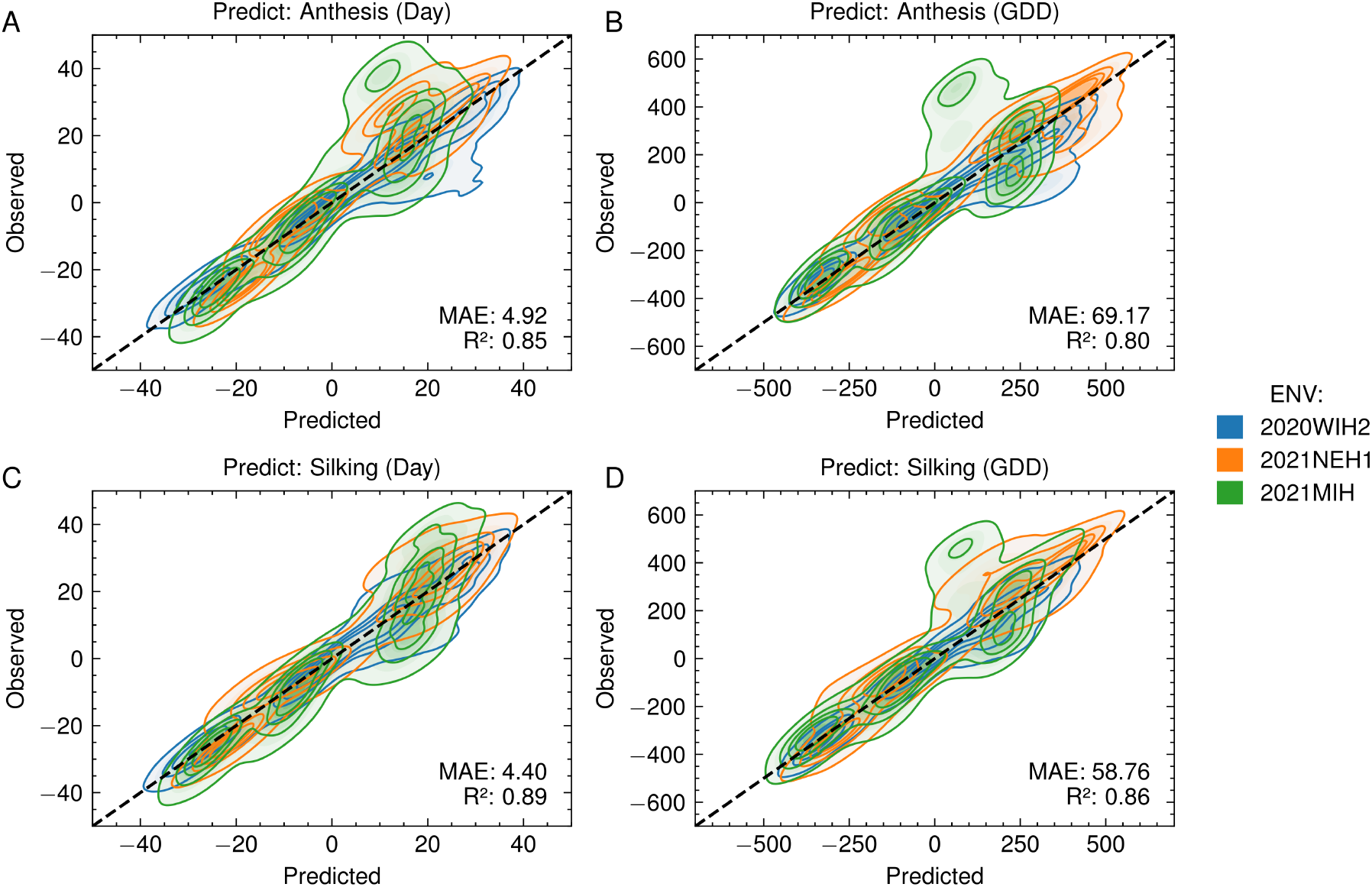
Predicted versus observed flowering times from models trained with UAV images captured within one month before or after flowering and labeled using the continuous-label framework. Predictions were made for anthesis (A, B) and silking (C, D) using either calendar days (A, C) or growing degree days (GDD; B, D). Each point represents an individual plot; colors indicate experimental locations: 2020WIH2 (blue), 2021NEH1 (orange), and 2021MIH (green). The dashed black line represents 1:1 correspondence. Mean absolute error (MAE) and coefficient of determination (*R*^2^) are shown in each panel.

**Figure 5.**
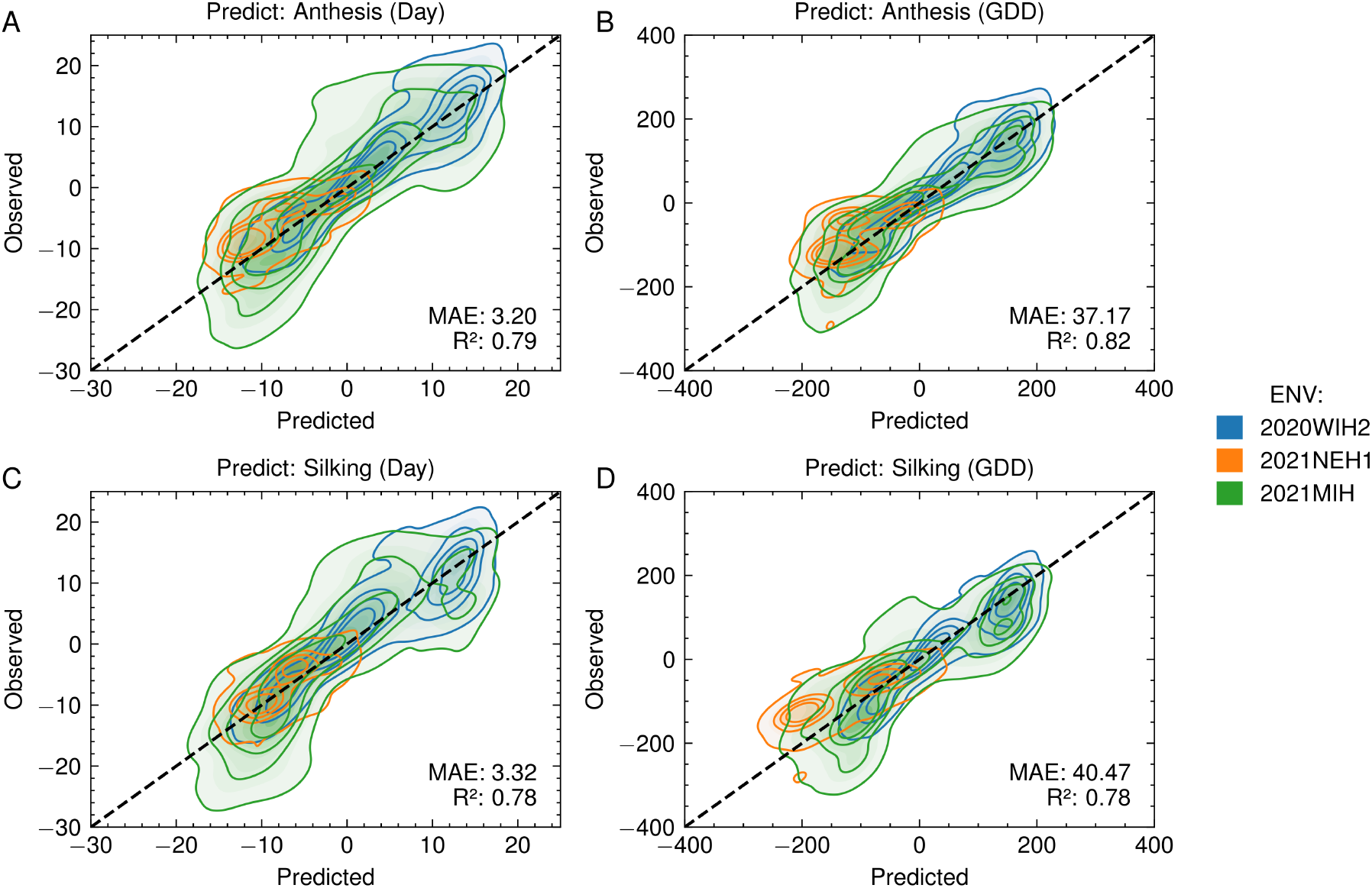
Predicted versus observed flowering times from models trained with UAV images captured within two weeks before or after flowering and labeled using the continuous-label framework. Predictions were made for anthesis (A, B) and silking (C, D) using either calendar days (A, C) or growing degree days (GDD; B, D). Each point represents an individual plot; colors indicate experimental locations: 2020WIH2 (blue), 2021NEH1 (orange), and 2021MIH (green). The dashed black line represents 1:1 correspondence. Mean absolute error (MAE) and coefficient of determination (*R*^2^) are shown in each panel.

To separate the effect of training-window selection from differences in evaluation range, we evaluated models trained using different temporal windows on identical held-out test subsets restricted to *±*2 weeks (a 28-day interval) and *±*1 week (a 14-day interval) around flowering. Because the *±*2-week training dataset was a subset of the *±*1-month (approximately 60-day interval) dataset, models trained using the two temporal windows could be compared directly on exactly the same test images. This allowed models trained using the *±*1-month and *±*2-week datasets to be compared on exactly the same test images. When evaluated on the same *±*2-week test subset, models trained on imagery collected within two weeks of flowering consistently outperformed those trained on imagery collected within one month before or after flowering for both calendar-day and GDD targets (Table 1). When using data from the two weeks before and after flowering for training, the best predictive performance—regardless of whether calendar days or GDD were used as the target—was achieved when predicting the period from one week before to one week after flowering using imagery from the same interval. Models trained using the difference in calendar days as the target performed best when trained on two-week datasets and evaluated using UAV imagery from within one week of flowering, yielding MAEs of 2.61 days for anthesis and 2.87 days for silking. When GDD difference was the target, the lowest MAEs were likewise obtained using the two-week training dataset and imagery collected within one week of flowering, reaching 33.92 GDD for anthesis and 37.92 GDD for silking. Compared with models that used calendar days, GDD-based models demonstrated slightly better performance.

**Table 1.**
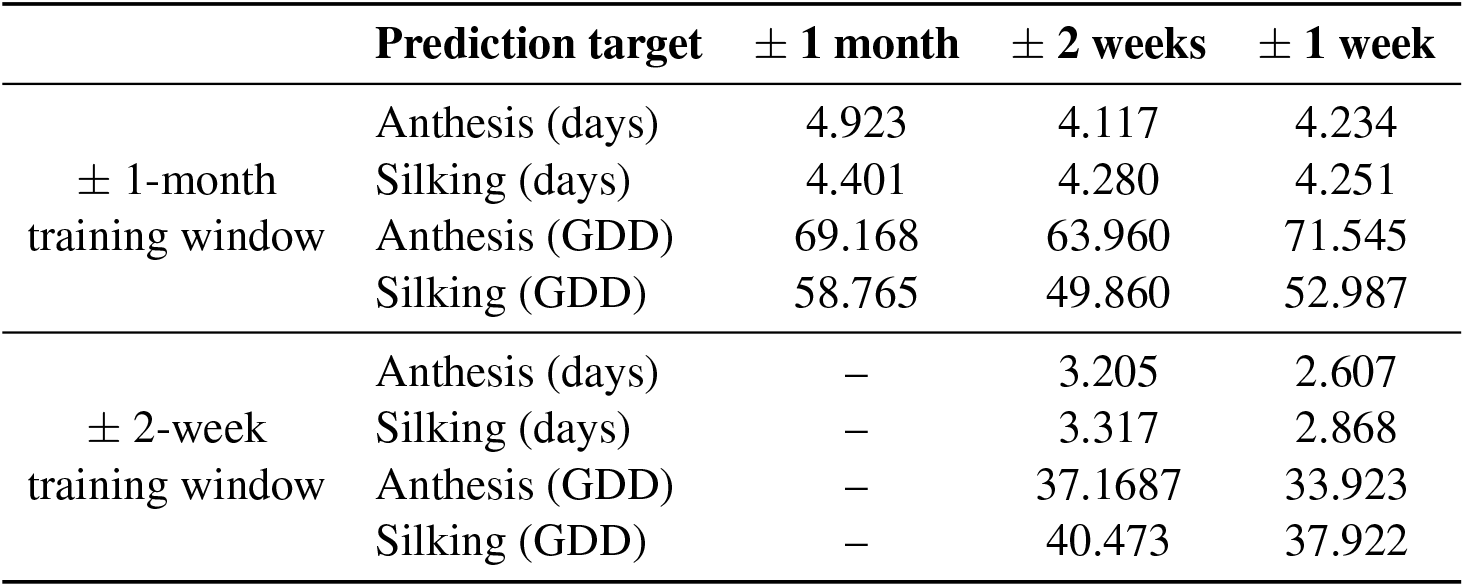
Mean absolute error by training and evaluation window.

| | Prediction target | $\pm 1$ month | $\pm 2$ weeks | $\pm 1$ week |
| --- | --- | --- | --- | --- |
| $\pm 1$-month<br>training window | Anthesis (days) | 4.923 | 4.117 | 4.234 |
|  | Silking (days) | 4.401 | 4.280 | 4.251 |
|  | Anthesis (GDD) | 69.168 | 63.960 | 71.545 |
|  | Silking (GDD) | 58.765 | 49.860 | 52.987 |
| $\pm 2$-week<br>training window | Anthesis (days) | – | 3.205 | 2.607 |
|  | Silking (days) | – | 3.317 | 2.868 |
|  | Anthesis (GDD) | – | 37.1687 | 33.923 |
|  | Silking (GDD) | – | 40.473 | 37.922 |

The three environments in the held-out test dataset were the 2020 Wisconsin G2F hybrid field 2, the 2021 Nebraska G2F hybrid field 1, and the 2021 Michigan G2F hybrid field. Prediction performance varied noticeably among these locations (Figure 6); 2020WIH2 consistently achieved the best predictive performance, whereas 2021MIH consistently exhibited the lowest performance across nearly all dataset configurations and temporal metrics. These locations differed in their environmental conditions and in the UAV platforms, cameras, and flight altitudes used to collect imagery. However, given the limited number of environments in the held-out dataset, it was not possible to determine which factors explained the substantial differences in model performance among locations. Averaging flowering-time estimates generated independently for the same plot from flights on different dates consistently decreased MAE, with diminishing improvements as the number of flights included in the average increased (Figure 7; Supplementary Figure S1). This pattern was consistent when considering flights collected within either two weeks or one month before or after flowering. Repeatability analysis showed that flowering-time estimates from UAV imagery could approach the repeatability of manual measurements, particularly when predictions from multiple flights were averaged (Figure 8). In 2020WIH2, repeatability increased as additional flights were included and, for several trait and prediction-scale combinations, approached the corresponding manual reference values. In 2021MIH, averaging across flights increased repeatability to levels that, for some trait and prediction-scale combinations, exceeded the manual reference. Repeatability in 2021NEH1 was generally lower. Overall, averaging independent predictions across UAV surveys improved the reliability of genotype-level flowering phenotypes and, under some conditions, produced repeatability comparable to or greater than that of manually scored flowering time. Because many genotypes in the held-out environments were represented in only one biological replicate, we additionally evaluated repeatability using the Cullis estimator, which showed a similar pattern S2.

**Figure 6.**
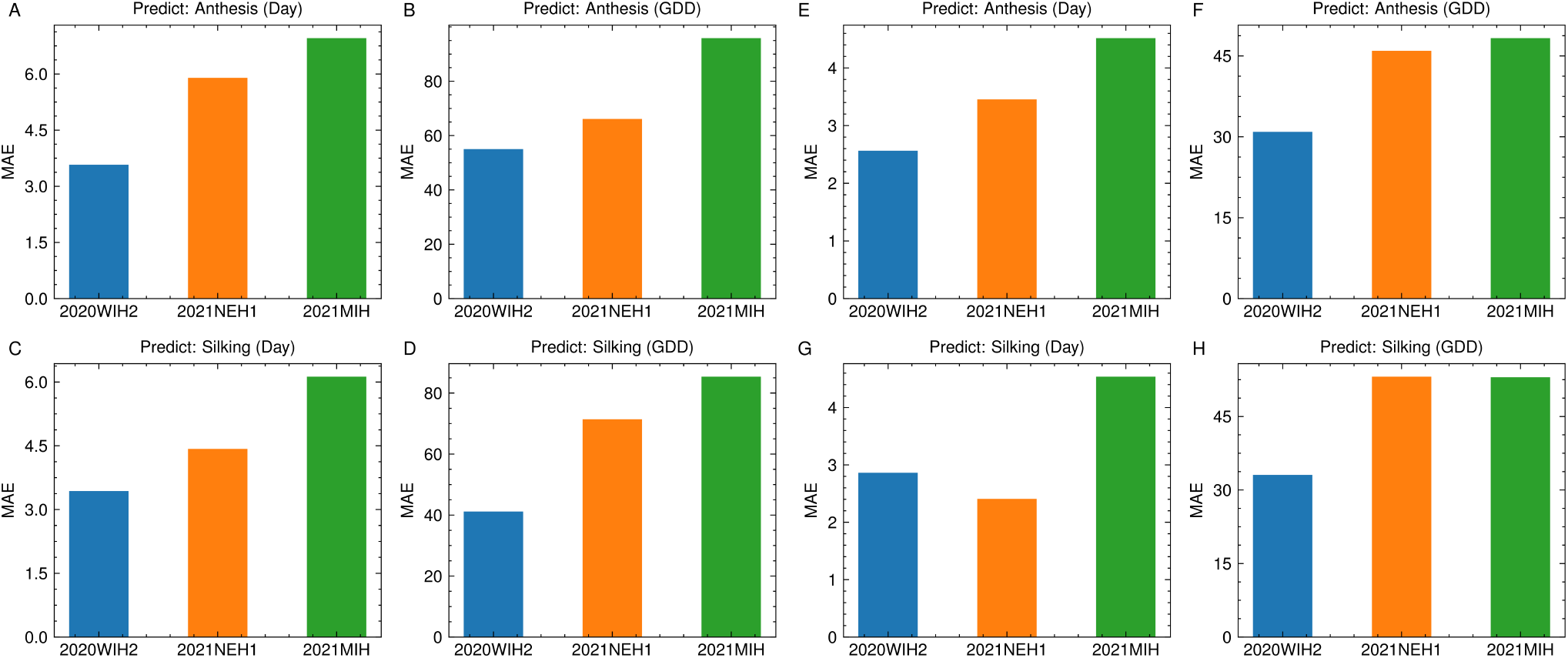
Mean absolute error (MAE) of flowering-time predictions across three experimental sites: 2020WIH2 (blue), 2021NEH1 (orange), and 2021MIH (green). Panels A–D show results from models trained with UAV images collected within one month before or after flowering, whereas panels E–H show results from models trained with images collected within two weeks before or after flowering. Anthesis predictions are shown in A, B, E, and F; silking predictions are shown in C, D, G, and H. Time units are calendar days (A, C, E, G) or growing degree days (GDD; B, D, F, H). All models were trained using the continuous-label framework.

**Figure 7.**
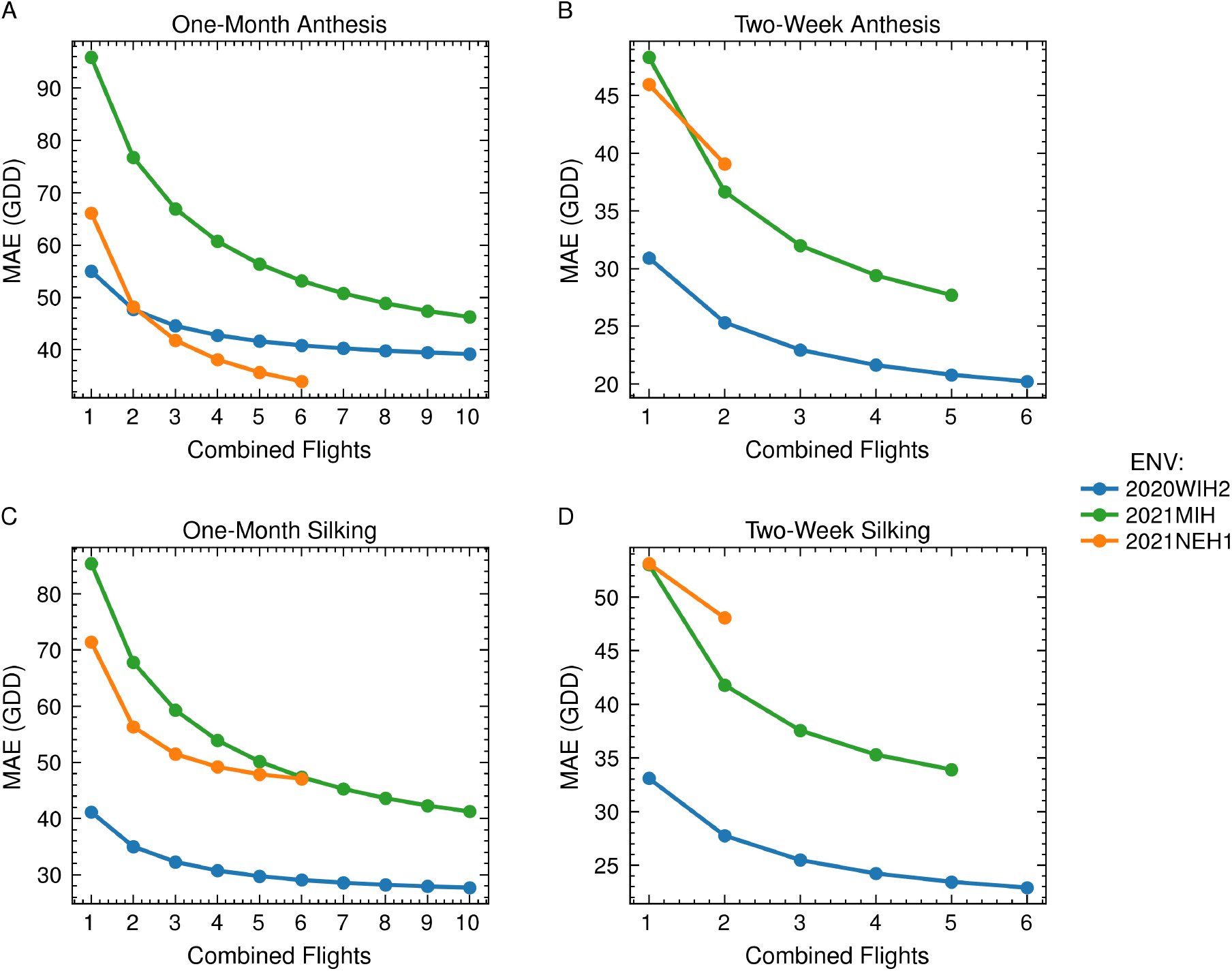
Effect of combining predictions from multiple UAV flights on flowering-time prediction error measured in growing degree days (GDD). Panels A and C show results from models trained with UAV images collected within one month before or after flowering, whereas panels B and D show results from models trained with images collected within two weeks before or after flowering. Anthesis predictions are shown in A and B, and silking predictions are shown in C and D. For each number of combined random flights, MAE was calculated across all available combinations containing that number of UAV flights.

**Figure 8.**
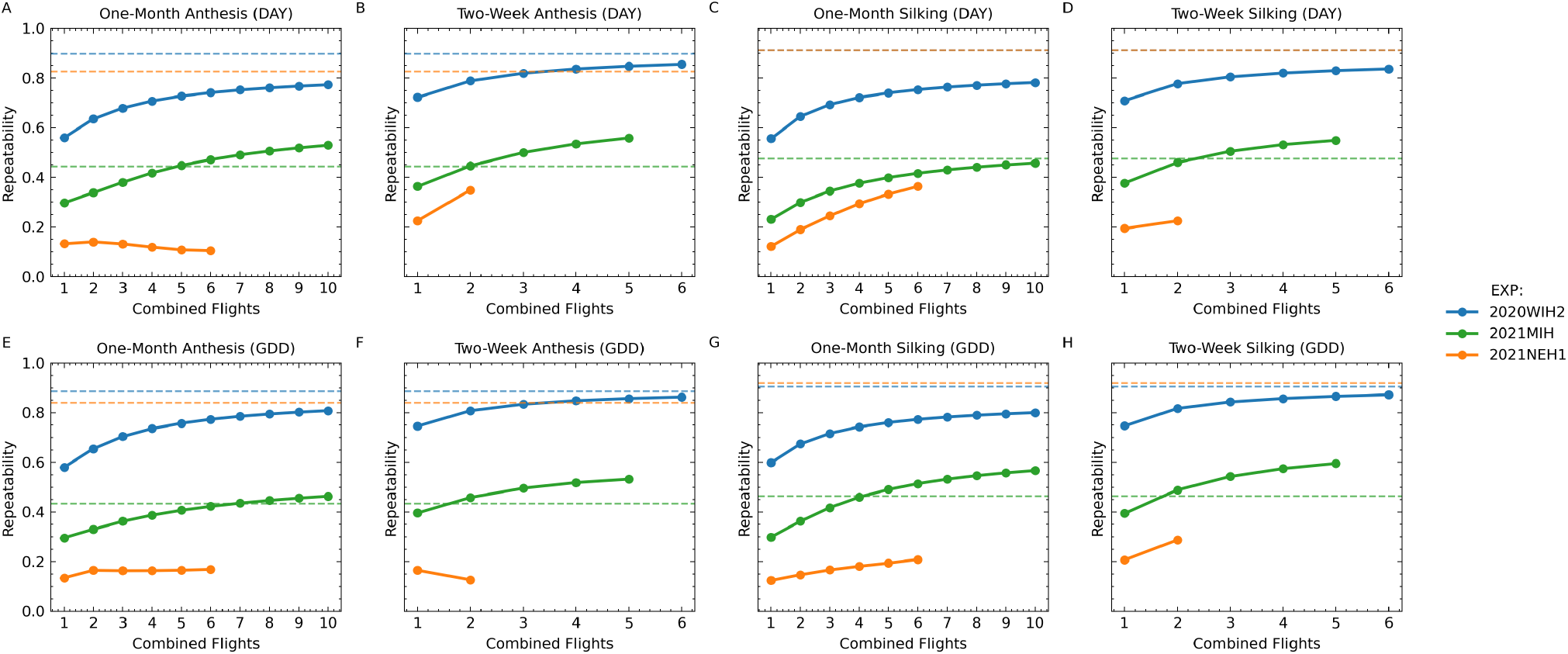
Effect of averaging flowering-time predictions from multiple UAV flights on repeatability. Panels A–D show repeatability for anthesis and silking predictions expressed in days (DAY), and panels E–H show the corresponding results expressed in growing degree days (GDD). One-month prediction windows are shown in A, C, E, and G, whereas two-week prediction windows are shown in B, D, F, and H. Solid lines show repeatability of UAV-derived flowering-time estimates for each held-out environment as predictions from increasing numbers of flights were averaged. Dashed horizontal lines indicate the repeatability of the corresponding manually scored flowering-time measurements. For each number of flights, repeatability was calculated for all available combinations containing that number of UAV flights and averaged across combinations.

Although the trained model outputs a predicted time relative to flowering, the same predictions can be used to classify individual plot images as flowered (positive predicted time relative to flowering) or not flowered (negative predicted time relative to flowering). In an independent dataset collected in Crawfordsville, Iowa, in 2022, for which observed flowering-time data were unavailable, the model’s flowering-status predictions agreed with human classifications of the same images in 92% of cases (Figure 9A). The images did not have sufficient resolution to reveal individual anthers, so human annotations of UAV images reflected the presence or absence of tassels (the male inflorescences of maize), which typically become visible several days before anthesis. Predicted flowering dates obtained from flights conducted 14 days apart were broadly consistent, with a mean absolute difference of 47 GDD (approximately 3 calendar days) between predictions for the same plot from the early and late flights (Figure 9B).

**Figure 9.**
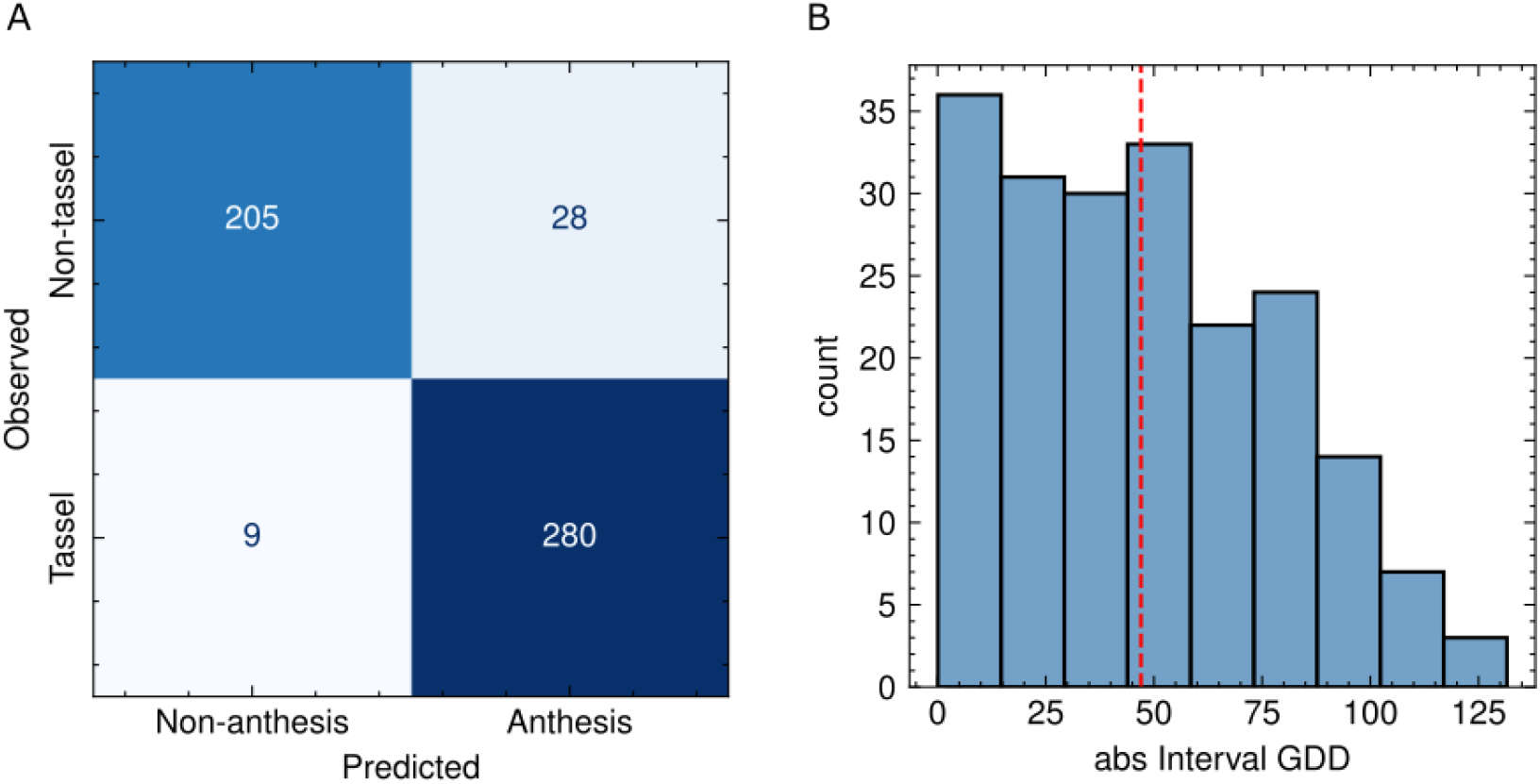
Evaluation of model performance on the 2022 Crawfordsville HIPS dataset. (A) Confusion matrix comparing model-predicted anthesis status (x-axis) with visual assessments of tassel presence (y-axis). Visual labels were assigned according to whether more than 50% of plants in a plot exhibited visible tassels. The model’s binary anthesis predictions aligned well with visually assessed tassel emergence. (B) Histogram of absolute differences in predicted anthesis GDD between two UAV flights for the same plots. Each bar represents the number of plots with a given level of disagreement between the two predictions. The red dashed line indicates the mean prediction difference and reflects model consistency across flights.

Model attention, as visualized by Grad-CAM, shifted dynamically across repeated observations of the same plot at different stages of maize development (Figure 10). Before visible tassel emergence, the model focused predominantly on the centers of leaf whorls, from which tassels would ultimately emerge. This pattern suggests that the model may recognize subtle pre-tasseling visual cues, such as upper-leaf orientation or stem extension. Around the anthesis date (i.e., within a few days before or after), highlighted areas were relatively small and less pronounced, suggesting more localized and uncertain attention. At substantially later stages, attention shifted from the tassel and plant center to the surrounding leaf canopy, which may be difficult to discern by human observers, suggesting that broader changes in canopy appearance became more informative for estimating how long before image collection flowering had occurred.

**Figure 10.**
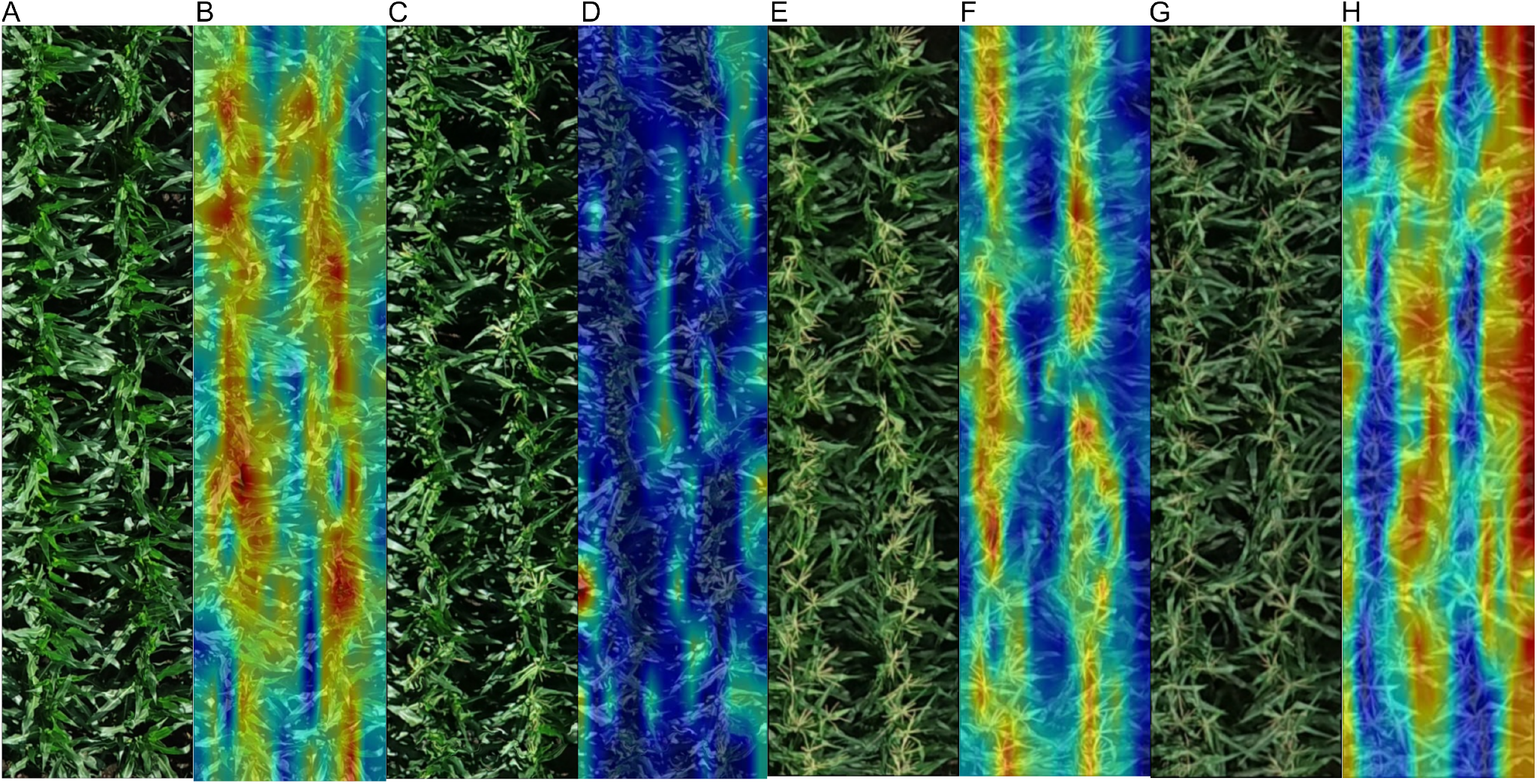
Orthomosaic images and corresponding Grad-CAM heat-map visualizations of a maize plot across four UAV survey dates. The plot ID is 2020WIH2:1020716, the genotype is W10004_0112/PHZ51, and recorded anthesis occurred on July 21. Images were generated by a two-week-window anthesis GDD prediction model. (A, B) The plot on July 12, with a reference label of *−*112.46 and a model prediction of *−*99.94. (C, D) The plot on July 20, with a label of *−*11.04 and a prediction of *−*17.12. (E, F) The plot on July 25, with a label of 46.98 and a prediction of 43.94. (G, H) The plot on August 5, with a label of 163.75 and a prediction of 201.40.

## DISCUSSION

The promise of high-throughput field phenotyping is to reduce the cost of trait measurement, enabling larger populations, greater replication, and broader evaluation across environments. However, because plant appearance, developmental timing, imaging conditions, and management practices can vary substantially among environments and years (Odé et al., 2025), many models remain closely tied to specific experimental settings or sampling schedules (Lyu et al., 2023; Camenzind and Yu, 2024; Fan et al., 2022). In this study, we developed a single regression framework using more than 200,000 plot-level UAV images from over 1,200 hybrid genotypes across 27 environments, spanning diverse years, locations, management conditions, imaging platforms, and flight schedules. By expressing flowering phenology as a continuous temporal prediction problem rather than a stage-specific classification task, data from heterogeneous experiments could be incorporated into a common modeling framework without requiring identical sampling schedules. This design emphasizes transferability across environments and better reflects how UAV phenotyping data are generated across breeding programs and research networks. More broadly, such an integrative framework may help move image-based phenotyping from experiment-specific models toward approaches that can be reused across diverse field conditions.

Models trained to predict flowering time via regression achieved reasonably strong performance. Using single UAV images collected within two weeks of flowering, models achieved MAEs of 3.20 and 3.32 days for anthesis and silking, respectively, across held-out environments; when GDD was used as the prediction target, MAEs were 37.17 GDD for anthesis and 40.47 GDD for silking (Figure 5). Combining predictions from multiple UAV surveys further reduced prediction error, with MAE decreasing substantially as additional surveys were included. These prediction errors are comparable to those of previous UAV-based approaches for estimating maize flowering phenology, which reported errors of approximately 3–6 days using multitemporal spectral, textural, or LiDAR-derived canopy-height information (Liu et al., 2025). In contrast, our approach relies only on standard RGB imagery and can operate with relatively sparse UAV surveys, substantially reducing data-acquisition requirements and the associated operational cost compared with approaches requiring specialized sensors or dense temporal sampling. Given that some locations in our dataset were flown only every 2–3 weeks, it was not possible to directly benchmark our model against classification-based methods. Such methods are not viable with the low-frequency UAV data used in this study and generated by many other research groups.

The repeatability analysis further suggests that UAV-derived flowering estimates can provide genotype-level phenotypes with reliability comparable to manual scoring, particularly when predictions from multiple flights are averaged. In 2021MIH, where flowering was relatively asynchronous among plants within the same plot, averaged multi-UAV predictions exceeded the repeatability of the manual measurements, suggesting that image-based estimates may be less sensitive to within-plot variation in flowering time.

One topic of ongoing debate in the research community is the degree of benefit, if any, provided by expressing the timing of crop developmental milestones in environmentally adjusted terms (e.g., GDD, photothermal time (Ellis et al., 1992), or heliothermal units (Zheng et al., 2023)) rather than simple calendar time. These adjusted measures are known to be more physiologically relevant, but they require more metadata and expertise to calculate than calendar-time intervals. Here, we found that the GDD-based MAEs corresponded to approximately 2.3 days for anthesis and 2.5 days for silking, compared with 3.20 and 3.32 days for models trained directly on calendar time. When flowering time can be expressed in GDD, it likely makes sense to do so; however, our results suggest that it is not essential, because prediction performance was only modestly reduced when models were trained and evaluated using calendar time. Although GDD accounts for temperature-driven differences in development, other environmental conditions may also affect the relationship between plant appearance and time relative to flowering. For pre-flowering predictions, environmental conditions between image acquisition and flowering have not yet occurred, whereas for retrospective predictions from post-flowering images, these conditions could potentially be reconstructed from historical environmental records. Future work could evaluate whether incorporating forecast, observed environmental, or thermal-time information as additional model inputs further improves prediction performance, although this would shift the task from image-based prediction toward a multimodal framework.

The ability to predict flowering time in advance or estimate it retrospectively substantially reduces the logistical burden of using UAVs to score this trait. However, prediction performance as flight dates moved farther from the recorded flowering time. Models trained using images collected within two weeks on either side of recorded flowering predicted flowering approximately one day more accurately than models trained using images collected within one month on either side of flowering (Table 1). The increase in MAE with broader prediction windows likely reflects the inclusion of observations that contain less direct information about flowering status. In the current framework, predictions from different flight dates are treated equally, whereas future approaches could incorporate temporal weighting or model uncertainty to give greater weight to observations collected closer to flowering. Flying more often can improve accuracy because averaging flowering-time estimates for the same plot imaged on different dates reduced error (Figure 7). However, most of the potential gain was achieved with two or three flights, with rapidly diminishing marginal benefits thereafter. In the held-out test environment where imagery was collected at the highest resolution (2020WIH2), MAE declined below two days once predictions from three flights were included. An MAE below two days may be suitable for many genetics and breeding applications, particularly when the objective is to obtain flowering-time estimates without repeated manual field visits. A portion of the remaining prediction error may also reflect human visual estimation error and temporal interpolation in manually recorded flowering dates, particularly when inclement weather or holidays caused larger gaps between ground-truth measurements. Collecting imagery from a given field once every 10–14 days would be sufficient to provide multiple images within two weeks of flowering. With larger and more standardized datasets, future studies could move beyond evaluating whether flowering time can be predicted from sparse UAV observations to identifying optimal data-collection strategies. For example, systematic comparisons of image resolution, flight timing relative to flowering, flight altitude, and the number and spacing of flights could help determine which combinations provide the greatest predictive value. Such analyses could ultimately guide the development of more efficient UAV phenotyping protocols that minimize data-collection costs while maintaining accurate flowering-time estimates. Importantly, the model was evaluated in environments that were completely excluded from training, indicating that application to a new environment does not inherently require local flowering-time labels for model training. The independent Crawfordsville dataset further showed that useful predictions could be obtained without plot-level field-recorded anthesis dates. For breeding programs seeking greater site-specific performance, a relatively small locally scored subset could potentially be used for calibration or further adaptation.

UAV surveys conducted near the flowering date tended to produce predicted values close to zero and minimal heat-map activation, likely because flowering was used as the temporal reference point for label assignment. However, before flowering, attention focused on the tassel region—even before its clear emergence—suggesting that the model can detect subtle pre-flowering traits, such as visual changes in upper-canopy morphology, that are often imperceptible to human observers. This finding implies a capacity to capture temporally relevant signals beyond obvious phenotypic markers. The later shift in attention toward surrounding leaf tissues highlights the model’s ability to reweight visual features depending on developmental context. This capacity suggests that the trained backbone can serve as a foundation for other developmental-stage recognition tasks, particularly those involving subtle phenotypic changes over time (Zdrazil et al., 2025).

## CONCLUSION

Regression-based approaches applied to UAV-based RGB imaging have strong potential to predict crop flowering time using data from a modest number of flights. The models evaluated here predicted flowering time with acceptable prediction performance in previously unseen environments using imagery from as few as one flight. Significant differences in prediction performance among locations suggest that site-specific factors, including environmental conditions, sensor quality, and image resolution, may influence model performance; further work is needed to identify optimal flight parameters and data-collection protocols. Examination of model interpretability suggests that the trained model learned key visual features and physiological transitions that precede and follow flowering, and that the trained backbone may be repurposed to quantify other developmental milestones under field conditions. The model’s performance in unseen environments suggests that it may already be useful to research groups seeking to estimate flowering time from previously collected UAV-based RGB imagery.

## DATA AVAILABILITY

The UAV imagery and associated spatial data used in this study were obtained from a combination of publicly available datasets and data generated by our group. Publicly available datasets were obtained through the Genomes to Fields (G2F) 2020-2021 project (DeSalvio et al., 2026) and the USDA/NIFA High-Intensity Phenotyping Sites (HIPS) program (Shrestha et al., 2024). Environment-specific G2F 2020-2021 UAV datasets are available through Figshare and hosted via Purdue University’s Data 2 Science (D2S) platform, while HIPS data are available through Dryad (Shrestha et al., 2024). UAV imagery and associated data generated by our group for the remaining environments are available through Zenodo. Repository information and DOIs for all environments used in this study are provided in Table S2. The Zenodo repository listed as “All other locations” in Table S2 (10.5281/zenodo.22802687) also contains the curated, analysis-ready HDF5 dataset used in this study, including plot-level UAV images matched with flowering-time phenotypes and associated metadata, as well as the trained model weights used for prediction. The source code used for data preprocessing, model training, prediction, evaluation, statistical analyses, and generation of the results presented in this study is publicly available through our GitHub repository.

## ACKNOWLEDGMENTS

This work was supported by the Nebraska Corn Board; the U.S. Department of Agriculture’s National Institute of Food and Agriculture under Award No. 2020-68013-30934; the National Science Foundation under Award No. IOS-2412928 to JCS; and a Heuermann Postdoctoral Fellowship to ZJ. G2F data collection used in this study was supported by the National Corn Growers Association, the Iowa Corn Promotion Board, the Nebraska Corn Board, the Corn Marketing Program of Michigan, the Texas Corn Producers Board, USDA–ARS, and USDA–NIFA Hatch funds. Data collection in Texas was additionally supported by USDA–NIFA–AFRI under Award Nos. 2020-68013-32371 and 2021-67013-33915. The processing and availability of the 2020–2021 G2F UAV datasets were supported in part by Agriculture Genome to Phenome Initiative (AG2PI) seed grant (2022-70412-38454). A.J.D. was supported by the National Science Foundation Graduate Research Fellowship Program (2023–2026).

## SUPPLEMENTARY INFORMATION

**Figure S1.**
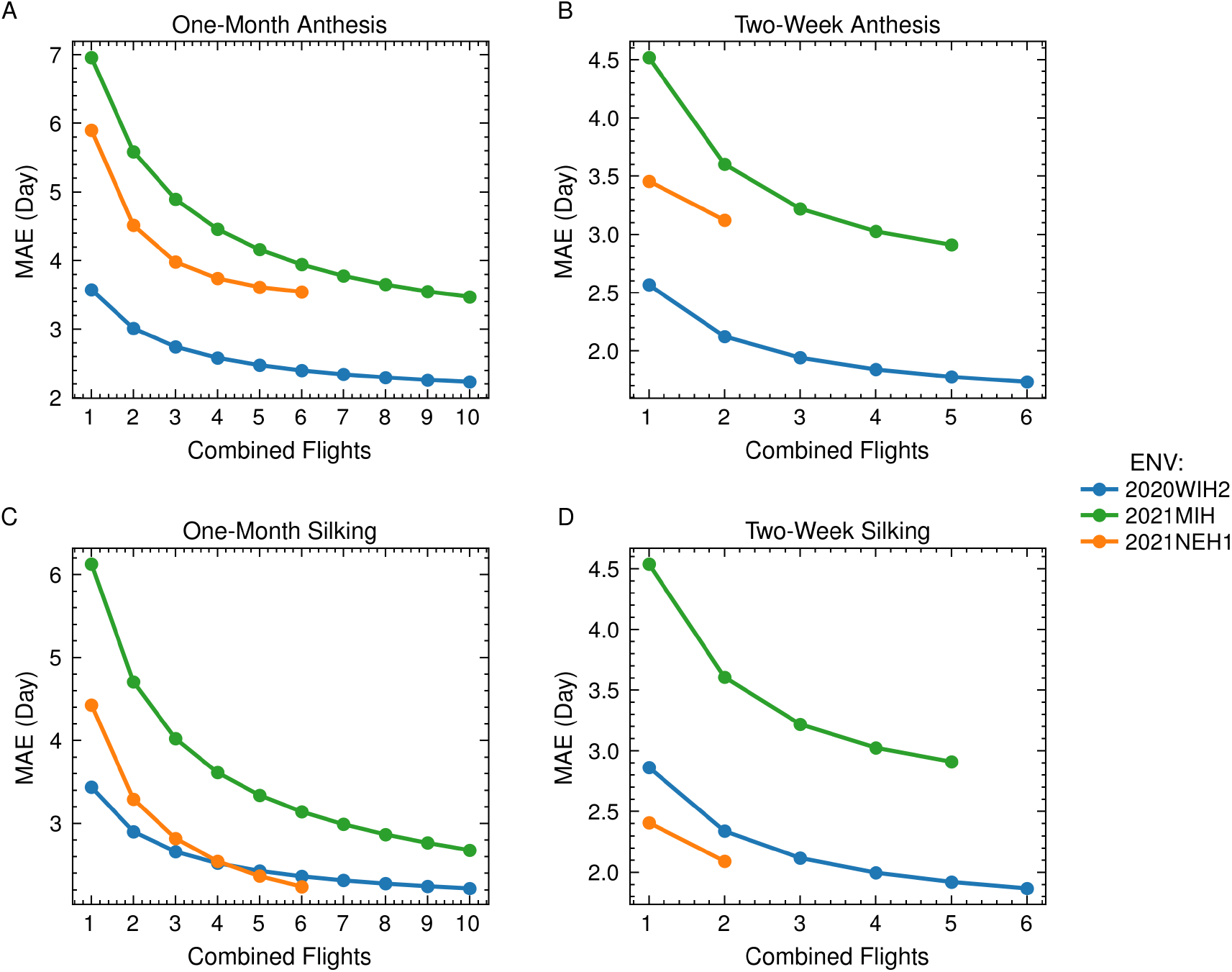
Effect of combining predictions from multiple UAV flights on flowering-time prediction error measured in Calendar days. Panels A and C show results from models trained with UAV images collected within one month before or after flowering, whereas panels B and D show results from models trained with images collected within two weeks before or after flowering. Anthesis predictions are shown in A and B, and silking predictions are shown in C and D. For each number of combined random flights, MAE was calculated across all available combinations containing that number of UAV flights.

**Table S1.** Number of UAV flights available within *±*1 month and *±*2 weeks of flowering for each environment.

| Environment | $\pm 1$ month | $\pm 2$ weeks | Environment | $\pm 1$ month | $\pm 2$ weeks |
| --- | --- | --- | --- | --- | --- |
| 2019MIH | 5 | 3 | 2021NEH1 | 6 | 2 |
| 2020DEH1 | 7 | 4 | 2021TXH1 | 12 | 8 |
| 2020MIH | 8 | 5 | 2021TXH2 | 12 | 7 |
| 2020MNH1 | 8 | 5 | 2021TXH3 | 15 | 7 |
| 2020MOH1 | 25 | 13 | 2021WIH1 | 13 | 8 |
| 2020TXH1 | 12 | 6 | 2021WIH2 | 13 | 7 |
| 2020TXH2 | 12 | 6 | 2021WIH3 | 12 | 7 |
| 2020TXH3 | 12 | 6 | 2022LIN | 3 | 2 |
| 2020WIH1 | 16 | 10 | 2022MIH | 9 | 6 |
| 2020WIH2 | 10 | 6 | 2022SCT | 3 | 2 |
| 2020WIH3 | 10 | 7 | 2023MIH | 6 | 2 |
| 2021IAH4 | 9 | 5 | 2023UNLHIPS | 5 | 3 |
| 2021MIH | 11 | 5 | 2024UNLG2F | 5 | 2 |
| 2021MNH1 | 6 | 4 | 2024UNLHIPS | 5 | 3 |

**Table S2.** Publicly available UAV datasets used in this study.

| Environment / Dataset | Repository | DOI |
| --- | --- | --- |
| 2020TXH1;2;3 | Figshare | 10.6084/m9.figshare.33301716 |
| 2021TXH1;2;3 | Figshare | 10.6084/m9.figshare.33301719 |
| 2021NEH1 | Figshare | 10.6084/m9.figshare.33301713 |
| 2021IAH4 | Figshare | 10.6084/m9.figshare.33301599 |
| 2020MOH1C5a | Figshare | 10.6084/m9.figshare.33301704 |
| 2020MOH1C5b | Figshare | 10.6084/m9.figshare.33301707 |
| 2020MNH1 | Figshare | 10.6084/m9.figshare.33301611 |
| 2021MNH1 | Figshare | 10.6084/m9.figshare.33301701 |
| 2020WIH1 | Figshare | 10.6084/m9.figshare.33301749 |
| 2020WIH2 | Figshare | 10.6084/m9.figshare.33438316 |
| 2020WIH3 | Figshare | 10.6084/m9.figshare.33301764 |
| 2021WIH1 | Figshare | 10.6084/m9.figshare.33301755 |
| 2021WIH2 | Figshare | 10.6084/m9.figshare.33301758 |
| 2021WIH3 | Figshare | 10.6084/m9.figshare.33301776 |
| HIPS multistate maize trials | Dryad | 10.5061/dryad.905qfttm |
| All other locations | Zenodo | 10.5281/zenodo.22802687 |

**Figure S2.**
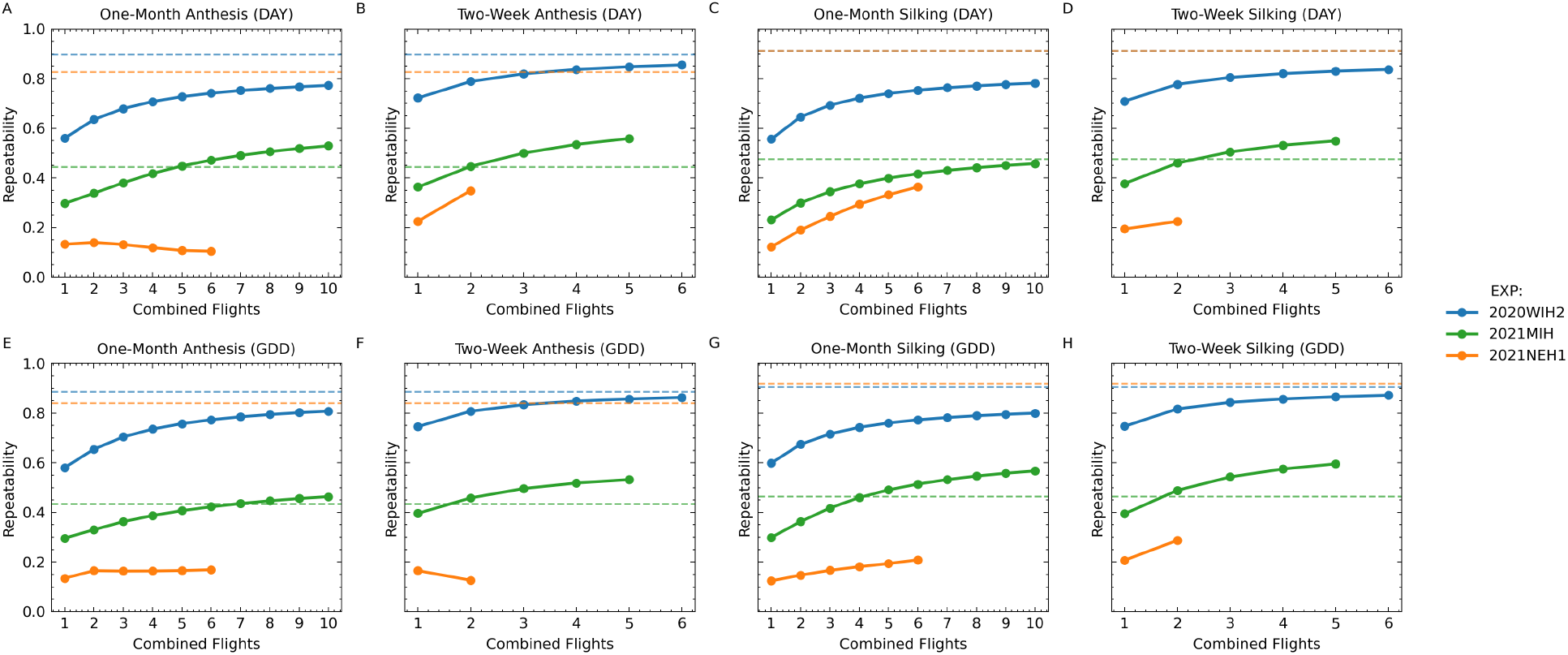
Effect of averaging flowering-time predictions from multiple UAV flights on Cullis repeatability. Panels A–D show Cullis repeatability for anthesis and silking predictions expressed in days (DAY), and panels E–H show the corresponding results expressed in growing degree days (GDD). One-month prediction windows are shown in A, C, E, and G, whereas two-week prediction windows are shown in B, D, F, and H. Solid lines show Cullis repeatability of UAV-derived flowering-time estimates for each held-out environment as predictions from increasing numbers of flights were averaged. Dashed horizontal lines indicate the Cullis repeatability of the corresponding manually scored flowering-time measurements. For each number of flights, repeatability was calculated for all available combinations containing that number of UAV flights and averaged across combinations.

## REFERENCES

Agisoft LLC (n.d.). Agisoft metashape. Photogrammetry software.

AlKhalifah, N., Campbell, D. A., Falcon, C. M., Gardiner, J. M., Miller, N. D., Romay, M. C., Walls, R., Walton, R., Yeh, C.-T., Bohn, M., et al. (2018). Maize genomes to fields: 2014 and 2015 field season genotype, phenotype, environment, and inbred ear image datasets. BMC research notes, 11(1):452.

Bonhomme, R., Derieux, M., and Edmeades, G. (1994). Flowering of diverse maize cultivars in relation to temperature and photoperiod in multilocation field trials. Crop science, 34(1):156–164.

Borrás, L., Astini, J. P., Westgate, M. E., and Severini, A. D. (2009). Modeling anthesis to silking in maize using a plant biomass framework. Crop Science, 49(3):937–948.

Buckler, E. S., Holland, J. B., Bradbury, P. J., Acharya, C. B., Brown, P. J., Browne, C., Ersoz, E., Flint-Garcia, S., Garcia, A., Glaubitz, J. C., et al. (2009). The genetic architecture of maize flowering time. Science, 325(5941):714–718.

Camenzind, M. P. and Yu, K. (2024). Multi temporal multispectral uav remote sensing allows for yield assessment across european wheat varieties already before flowering. Frontiers in plant science, 14:1214931.

Chang, A., Yeom, J., Jung, J., and Landivar, J. (2020). Comparison of canopy shape and vegetation indices of citrus trees derived from uav multispectral images for characterization of citrus greening disease. Remote Sensing, 12(24):4122.

Cross, H. and Zuber, M. (1972). Prediction of flowering dates in maize based on different methods of estimating thermal units 1. Agronomy journal, 64(3):351–355.

Cross, H. and Zuber, M. (1973). Interrelationships among plant height, number of leaves, and flowering dates in maize 1. Agronomy Journal, 65(1):71–74.

Cui, M., Jia, B., Liu, H., Kan, X., Zhang, Y., Zhou, R., Li, Z., Yang, L., Deng, D., and Yin, Z. (2017). Genetic mapping of the leaf number above the primary ear and its relationship with plant height and flowering time in maize. Frontiers in Plant Science, 8:1437.

DeSalvio, A. J., Mohseni, P., Adak, A., Murray, S. C., Arik, M. A., Wong, R. K. W., Jung, J., Lima, D. C., Aviles, A. C., Buckler, E. S., Duffield, N., Edwards, J., Ertl, D., Flint-Garcia, S., Gore, M. A., Hirsch, C. N., Holland, J. B., Kaeppler, S. M., Miller, J., Romay, C., Schnable, J. C., Singh, M. P., Sparks, E. E., Thompson, A., Washburn, J. D., Weldekidan, T., Winans, N. D., and de Leon, N. (2026). An open field phenomics resource for multimodal maize yield prediction across divergent environments. bioRxiv.

Deva, C., Dixon, L., Urban, M., Ramirez-Villegas, J., Droutsas, I., and Challinor, A. (2024). A new framework for predicting and understanding flowering time for crop breeding. Plants, People, Planet, 6(1):197–209.

Dosovitskiy, A., Beyer, L., Kolesnikov, A., Weissenborn, D., Zhai, X., Unterthiner, T., Dehghani, M., Minderer, M., Heigold, G., Gelly, S., et al. (2020). An image is worth 16×16 words: Transformers for image recognition at scale. arXiv preprint arXiv:2010.11929.

Ellis, R., Summerfield, R., Edmeades, G., and Roberts, E. (1992). Photoperiod, temperature, and the interval from sowing to tassel initiation in diverse cultivars of maize. Crop Science, 32(5):1225–1232.

Fan, J., Zhou, J., Wang, B., de Leon, N., Kaeppler, S. M., Lima, D. C., and Zhang, Z. (2022). Estimation of maize yield and flowering time using multi-temporal uav-based hyperspectral data. Remote Sensing, 14(13):3052.

Gilmore Jr, E. and Rogers, J. (1958). Heat units as a method of measuring maturity in corn 1. Agronomy journal, 50(10):611–615.

Ji, Z.-j., Ge, Y.-f., and Schnable, J. C. (2025). Scalable methods for quantifying the stay green ability of corn for yield prediction by using satellite image. agriRxiv.

Jin, H. and Eklundh, L. (2014). A physically based vegetation index for improved monitoring of plant phenology. Remote Sensing of Environment, 152:512–525.

Li, D., Wang, X., Zhang, X., Chen, Q., Xu, G., Xu, D., Wang, C., Liang, Y., Wu, L., Huang, C., et al. (2016). The genetic architecture of leaf number and its genetic relationship to flowering time in maize. New Phytologist, 210(1):256–268.

Li, J., Wang, E., Qiao, J., Li, Y., Li, L., Yao, J., and Liao, G. (2023). Automatic rape flower cluster counting method based on low-cost labelling and uav-rgb images. Plant Methods, 19(1):40.

Lima, D. C., Aviles, A. C., Alpers, R. T., Perkins, A., Schoemaker, D. L., Costa, M., Michel, K. J., Kaeppler, S., Ertl, D., Romay, M. C., et al. (2023). 2020-2021 field seasons of maize gxe project within the genomes to fields initiative. BMC research notes, 16(1):219.

Liu, Y., Nie, C., Li, L., Shi, L., Liu, S., Nan, F., Cheng, M., Yu, X., Bai, Y., Jia, X., et al. (2025). Maize tasseling date forecast from canopy height time series estimated by uav lidar data. The Crop Journal, 13(3):975–990.

Lyu, M., Lu, X., Shen, Y., Tan, Y., Wan, L., Shu, Q., He, Y., He, Y., and Cen, H. (2023). Uav time-series imagery with novel machine learning to estimate heading dates of rice accessions for breeding. Agricultural and Forest Meteorology, 341:109646.

Odé, A., Smith, N., Rebel, K., and de Boer, H. (2025). Temporal constraints on leaf-level trait plasticity for next-generation land surface models. Annals of Botany, page mcaf045.

Pauli, D., Chapman, S. C., Bart, R., Topp, C. N., Lawrence-Dill, C. J., Poland, J., and Gore, M. A. (2016). The quest for understanding phenotypic variation via integrated approaches in the field environment. Plant Physiology, 172(2):622–634.

Pix4D SA (n.d.). Pix4dmapper. Photogrammetry software.

Roberts, D. R., Bahn, V., Ciuti, S., Boyce, M. S., Elith, J., Guillera-Arroita, G., Hauenstein, S., Lahoz-Monfort, J. J., Schröder, B., Thuiller, W., et al. (2017). Cross-validation strategies for data with temporal, spatial, hierarchical, or phylogenetic structure. Ecography, 40(8):913–929.

Selvaraju, R. R., Cogswell, M., Das, A., Vedantam, R., Parikh, D., and Batra, D. (2017). Grad-cam: Visual explanations from deep networks via gradient-based localization. In Proceedings of the IEEE international conference on computer vision, pages 618–626.

Shepard, N. R., DeSalvio, A. J., Arik, M., Adak, A., Murray, S. C., and Varela, J. I. (2025). Deep learning-based high-throughput detection of flowered maize (zea mays l.) plots from uas imagery across environments. The Plant Phenome Journal, 8(1):e70021.

Shrestha, N., Powadi, A., Davis, J., et al. (2024). Crop performance, aerial, and satellite data from multistate maize yield trials. Dataset.

Srivastava, N., Hinton, G., Krizhevsky, A., Sutskever, I., and Salakhutdinov, R. (2014). Dropout: a simple way to prevent neural networks from overfitting. The journal of machine learning research, 15(1):1929–1958.

Thornton, M. and Devarakonda, R. (2024). Daymet single pixel extraction tool.

Tollenaar, M., Daynard, T., and Hunter, R. (1979). Effect of temperature on rate of leaf appearance and flowering date in maize 1. Crop Science, 19(3):363–366.

Vanbrabant, Y., Delalieux, S., Tits, L., Pauly, K., Vandermaesen, J., and Somers, B. (2020). Pear flower cluster quantification using rgb drone imagery. Agronomy, 10(3):407.

Vidican, R., Mălinaş, A., Ranta, O., Moldovan, C., Marian, O., Gheţe, A., Ghişe, C. R., Popovici, F., and Cătunescu, G. M. (2023). Using remote sensing vegetation indices for the discrimination and monitoring of agricultural crops: A critical review. Agronomy, 13(12):3040.

Weis, A. E., Wadgymar, S. M., Sekor, M., and Franks, S. J. (2014). The shape of selection: using alternative fitness functions to test predictions for selection on flowering time. Evolutionary ecology, 28(5):885–904.

Žalud, Z., Hlavinka, P., Prokeš, K., Semerádová, D., Jan, B., and Trnka, M. (2017). Impacts of water availability and drought on maize yield–a comparison of 16 indicators. Agricultural Water Management, 188:126–135.

Zdrazil, J., Kong, L., Klimeš, P., Jasso-Robles, F. I., Saiz-Fernández, I., Güder, F., Spíchal, L., Snášel, V., and De Diego, N. (2025). Next-generation high-throughput phenotyping with trait prediction through adaptable multi-task computational intelligence. Computers and Electronics in Agriculture, 235:110390.

Zhao, B., Khound, R., Ghimire, D., Zhou, Y., Maharjan, B., Santra, D. K., and Shi, Y. (2022). Heading percentage estimation in proso millet (panicum miliaceum l.) using aerial imagery and deep learning. The plant phenome journal, 5(1):e20049.

Zheng, J., Zhang, F.-c., et al. (2023). Novel models for simulating maize growth based on thermal time and photothermal units: Applications under various mulching practices. Journal of Integrative Agriculture, 22(5):1381–1395.

Zhuang, F., Qi, Z., Duan, K., Xi, D., Zhu, Y., Zhu, H., Xiong, H., and He, Q. (2020). A comprehensive survey on transfer learning. Proceedings of the IEEE, 109(1):43–76.

